# A transcriptome-wide meta-analysis reveals lack of cancer-cell intrinsic determinants of response to immune checkpoint blockade

**DOI:** 10.1101/2023.07.30.551135

**Authors:** Yu Amanda Guo, Tanmay Kulshrestha, Mei Mei Chang, Irfahan Kassam, Egor Revkov, Simone Rizzetto, Aaron C. Tan, Daniel S.W. Tan, Iain Beehuat Tan, Anders Jacobsen Skanderup

## Abstract

Immune-checkpoint therapy (ICB) has conferred significant and durable clinical benefit to some cancer patients. However, most patients do not respond to ICB, and reliable biomarkers of ICB response are needed to improve patient stratification. Here, we performed a transcriptome-wide meta-analysis across 1,486 tumors from ICB-treated patients and tumors with expected ICB outcomes based on microsatellite status. Using a robust transcriptome deconvolution approach, we inferred cancer and stroma-specific gene expression differences and identified cell-type specific features of ICB response across cancer types. Consistent with current knowledge, stromal expression of *CXCL9*, *CXCL13*, and *IFNG* were the top determinants of favorable ICB response. In addition, we identified a group of potential immune-suppressive genes, including *FCER1A*, associated with poor response to ICB. Strikingly, PD-L1 expression in stromal cells, but not cancer cells, is correlated with ICB response across cancer types. Furthermore, the unbiased transcriptome-wide analysis failed to identify cancer-cell intrinsic features of ICB response conserved across tumor types. Overall, our results challenge the prevailing dogma that cancer cells present tissue-agnostic molecular markers that modulate immune activity and ICB treatment response. These results have implications for the development of improved ICB treatments and diagnostics.

## Introduction

Immune checkpoint blockade therapy (ICB) is an established class of immunotherapy that induces significant tumor shrinkage and long-term disease control in many cancers. However, only a subset of patients benefit from ICB, with response rates ranging from 0-3% in pancreatic cancer^1^ to about 40% in melanoma and microsatellite instable (MSI) tumors^2^. Presently, the Food and Drug Administration (FDA)-approved biomarkers for ICB are PD-L1 expression^3^, microsatellite instability^4^, and high tumor mutation burden (TMB) of >10mutations/Mb^5^. While PD-L1 expression is an obvious biomarker for PD-1/PD-L1 blockade therapies, it is only weakly predictive of treatment response in many solid cancer types^6^. TMB is a measure of tumor immunogenicity, where increased mutation load is associated with higher neo-antigen burden and anti-tumoral immune responses. However, the predictive power of TMB varies significantly by cancer type and it is difficult to define a single TMB threshold across cancer types^7,8^. Likewise, MSI tumors are more immunogenic, as defective mismatch repair results in large numbers of insertions and deletions in the cancer genomes. In practice, MSI status has limited applications as a biomarker of ICB response, since only a small subset of solid tumors are microsatellite instable^9,10^.

Transcriptional signatures of tumor immune phenotypes have been proposed to predict for ICB response^11–13^. However, analysis of large-scale ICB trial data found that existing transcriptional signatures often have limited predictive accuracy in new patient cohorts^6^. Two recent meta-analyses investigated the genomic and transcriptomic predictors of ICB response^14,15^. Litchfield et al.^14^ evaluated previously described biomarkers of ICB response, and found TMB and *CXCL9* expression as the most consistent predictors of ICB response. Bareche et al. performed a comparative analysis of existing gene signatures of ICB response, and proposed a new predictive gene signature ^15^. However, existing large-scale studies have been restricted to bulk tumor transcriptomic data, providing limited insights into the roles of cancer and stroma cells in the tumor microenvironment (TME) during ICB therapy.

Activation of effector immune cells such as T-cells and natural killer (NK) cells are dependent on a balance of co-stimulatory and co-inhibitory interactions between these effector cells, antigen-presenting immune cells such as dendritic cells and macrophages, as well as cancer cells in the TME. Dysregulation of these ligand-receptor interactions, or immune checkpoints, leads to immunosuppression in the tumor. However, it is challenging to tease apart these complex interactions between cancer and immune cells using bulk tumor transcriptomics. For example, although inhibitory immune checkpoint ligand PD-L1 is a known biomarker of ICB response, it is unclear if cancer cells escape immune cell killing by expressing PD-L1, or recruiting PD-L1 immune cells, with studies supporting both scenarios^16^. Therefore, learning the cell type specificities of ICB biomarkers can provide insights on the underlying mechanisms of immune evasion and ICB resistance. While single- cell transcriptomics has been applied to explore immune cell types and molecular mechanisms associated with ICB response, single cell transcriptomics studies are currently limited to small patient cohorts^17,18^. The small sample size, coupled with high intra- and inter- patient heterogeneity and intrinsic noise in single-cell data, limits the power of single- cell transcriptomics for systematic biomarker discovery.

To study features of ICB response in cancer cells as well as in their neighboring stromal cells, we applied a transcriptome deconvolution technique to estimate stroma- and cancer-cell-specific expression of individual genes from bulk tumor transcriptomes. We used a robust and validated non-negative least squares approach^19^, which have previously been applied across 8,000 TCGA tumors from 20 solid cancer types to uncover ligand-receptor crosstalk and metabolic states of cancer and stroma cells in the TME^20 21^. Here, we applied this transcriptome deconvolution technique to study the differential expression signatures of ICB responders and non-responders in cancer and stroma cells. Using a cohort of 1,486 tumors with clinically annotated ICB response and tumors with expected ICB outcomes based on microsatellite status., our analysis revealed a lack of cancer-cell intrinsic gene expression signatures associated with ICB response across tumor types. In contrast, in stromal cells, we identified multiple immunomodulators and checkpoint genes recurrently associated with ICB response across cohorts and tumor types. Many of these genes have previously been linked with ICB response and we uncovered a subset of novel and potentially immune-suppressive genes negatively correlated with response. Using these conserved stromal-specific gene signatures, we developed a compact 3-feature predictive model of ICB response that demonstrated improved accuracy over existing biomarkers in unseen patient cohorts and tumor types.

## Results

### Patient cohorts used for transcriptome-wide analysis of ICB response

To systematically explore transcriptome-wide biomarkers of ICB response, we obtained transcriptome data for 534 tumors, comprising 4 ICB-treated patient cohorts from 3 tumor types (gastric cancer^22^, urothelial cancer^23^ and melanoma^24,25^; **Figure 1**, **Supplementary Table 1,** and **Methods**). Since about half of MSI tumors are expected to respond to ICB, as compared to <10% in microsatellite stable (MSS) tumors^2,26,27^, we further supplemented the discovery cohort using large cohorts of treatment-naïve MSI/MSS tumors. We obtained the transcriptomic profiles of 952 tumors from 3 tumor types in TCGA with the highest frequency of MSI tumors (colorectal cancer, gastric cancer and endometrial cancer, **Figure 1**). As Epstein–Barr virus (EBV)-positive gastric tumors are also enriched for ICB responders^22,27^, we grouped MSI and EBV tumors together as the responder-enriched group for gastric cancer. In total, the discovery cohort comprised of 1,486 tumors from 5 cancer types.

**Figure 1.**
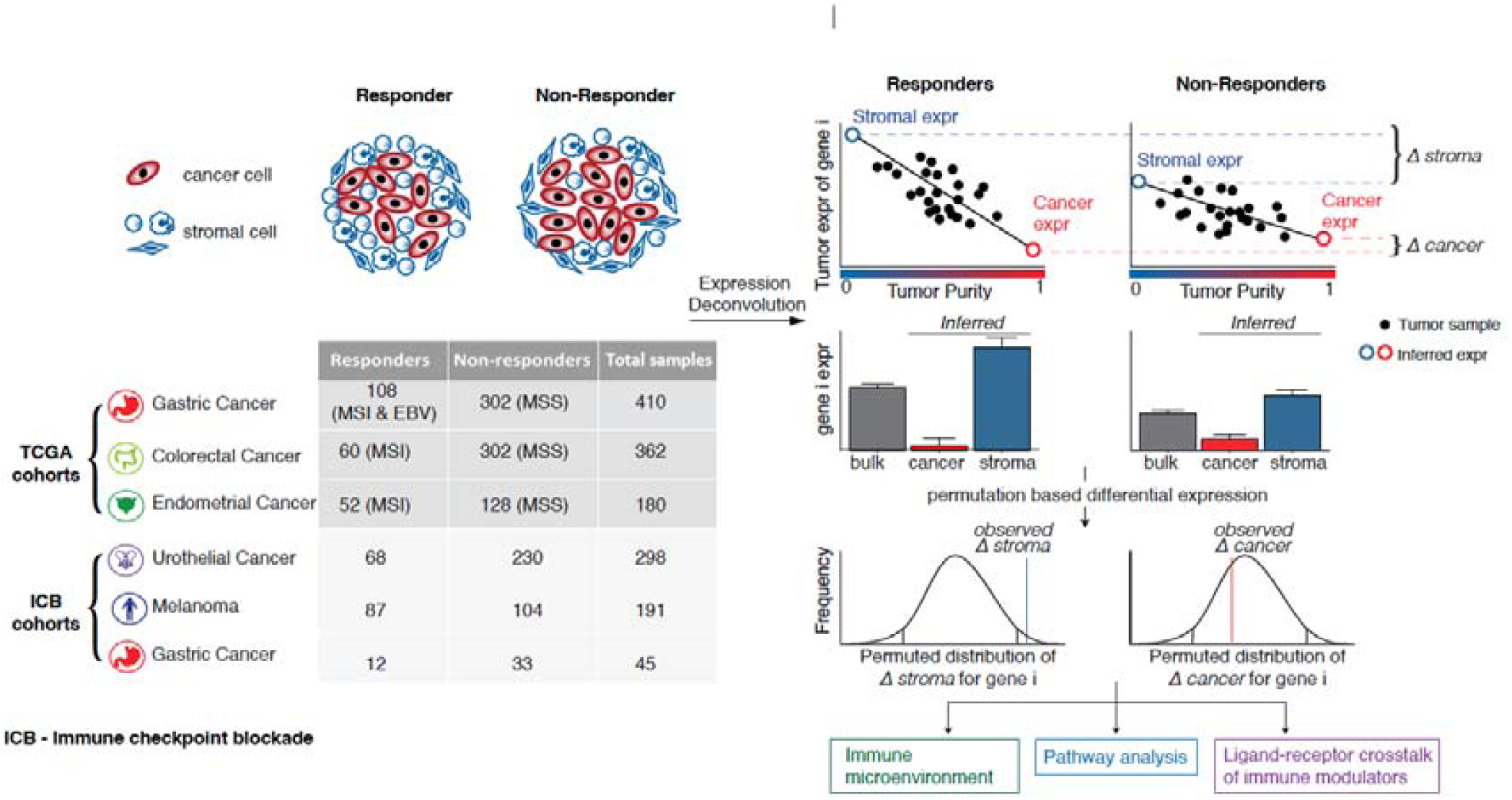
Data summary and schematic of workflow for the analysis of determinants of immunotherapy response.

### Tumor transcriptome deconvolution to identify features of ICB response in cancer and stromal cells

We performed tumor transcriptome deconvolution to estimate the stromal- and cancer-cell expression of each gene in tumors from responders (MSI) and non-responders (MSS) (**Figure 1**). Briefly, we first used a consensus-approach to infer tumor purities based on transcriptomic and genomic data (where available, Methods). For each ICB response group, we estimated gene expression levels in stromal and cancer cells using a non-negative least squares regression approach^20,21^. Finally, we identified cancer/stromal-cell differentially expressed genes (DEGs) between ICB responders and non-responders using a permutation-based statistic (**Figure 1, Supplementary Figure 1**).

### Top stromal genes associated with ICB response are enriched in immune pathways

From this unbiased transcriptome-wide analysis, we identified 59 genes with differential expression in stromal cells across all cohorts and tumor types (**Figure 2A-B, Supplementary Figure 2A and 2C, Methods**). Among these DEGs, 51 (86%) had immune-related functions as defined by the Gene Ontology (GO; see Methods), significantly higher than random expectation (*p* < 2.2e-16, Fisher’s Exact test). Stromal DEGs include genes involved in: immune checkpoint signaling (e.g. *CD274*, *LAG3*, *CTLA4*), interferon gamma signaling (e.g. *IFNG*, *IRF1*, *STAT1*), T-cell effector function (e.g. *PRF1*, *GZMA*, GZMB), and NK-cell mediated cytotoxicity (*KLRD1*, *KLRC2*). Next, we performed pathway analysis to quantify the enrichment of specific biological functions among the stromal DEGs associated with ICB response. We identified 31 enriched GO biological process terms, all of which were immune related (**Figure 2C**). These 31 terms could be further grouped into 4 clusters based on similarity (see Methods): T cell activation, chemokine signaling, interferon-gamma response, and natural killer cell mediated cytotoxicity **(Figure 2D**).

**Figure 2.**
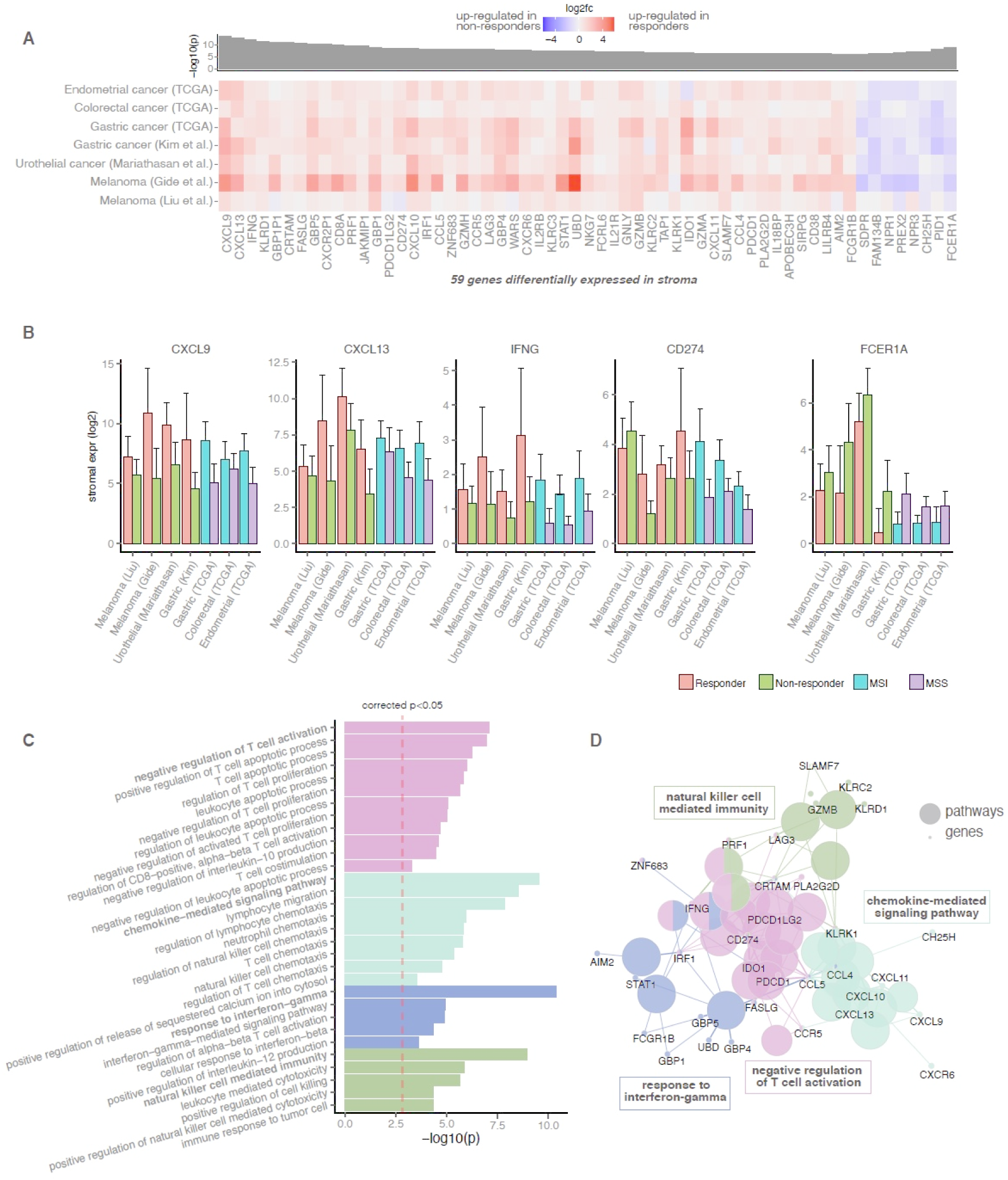
Differentially expressed genes and pathways between responders and non-responders in the stroma. **(A)** Heatmap of genes consistently differentially expressed across discovery cohorts in the stroma. Heatmaps are colored by the log2 fold change of gene expression. *P*-values of individual cohorts are combined using the Fisher’s method, and the -log10(meta p-value) is presented in the bar plot. Genes with q-value<0.01 and *p*- value<0.1 at least 1 ICB cohort are shown. **(B)** Bar plots of deconvoluted stromal expression of the top differentially expressed genes. Error bars represent estimated standard error. **(C)** Pathway enrichment of differentially expressed genes in the stroma. Bar plot shows the - log10(p-value) of significantly enriched pathways. Similar pathways are grouped by color. (D) Network representation of enriched pathways. Large nodes represent enriched pathways. Pathways with high similarity are fused and shown in the same color. Small nodes represent genes associated with each pathway.

The top-2 DEGs with the greatest overexpression in ICB responders were chemokines *CXCL9* and *CXCL13*. *CXCL9* is responsible for T-cell trafficking through binding the *CXCR3* receptor on T-cells and has been associated with increased T-cell infiltration in tumors^28^. *CXCL13* is involved in both B-cell and T-cell migration via binding to the *CXCR5* receptor. Production of *CXCL13* by T follicular helper cells (T_FH_) is essential for tertiary lymphoid structure formation at tumor sites as well as germinal center B cell activation^29^, which has been linked with anti-tumoral response to ICB^30^. Consistent with our meta-analysis, recent studies have also proposed *CXCL9* and *CXCL13* as key predictors of improved ICB response^31,32^. Remarkably, these two chemokines were also highlighted by a recent meta-analysis that evaluated existing biomarkers of ICB response^14^, suggesting that *CXCL9* and *CXCL13* are indeed the strongest individual transcriptomic predictors of ICB response independent of tumor type.

### Stromal genes negatively associated with ICB response

Although high immune infiltration is generally associated with better ICB response^11,33,34^, our analysis uncovered a group of immune-related genes consistently associated with poor ICB outcomes across tumor types. Among these genes, *FCER1A* showed the strongest and most consistent overexpression in ICB non-responders. *FCER1A* encodes a subunit of the IgE receptor. While *FCER1A* is an established marker of basophils, mast cells and dendritic cells, it is also expressed on immunosuppressive M2 macrophages^35^ and tumor associated macrophages (TAMs)^36^. *CH25H*, a gene encoding enzyme cholesterol 25-hydroxylase that catalyzes the formation of 25-hydroxycholesterol (25-HC), was also up-regulated in tumors of ICB non-responders. Interestingly, 25-HC is produced by macrophages in response to type-I interferon signalling and has numerous immunological effects including B-cell chemotaxis, macrophage differentiation, and regulation of inflammatory response^37^. We also observed two natriuretic peptides, NPR1 and NPR3, among genes up-regulated in ICB non-responders. Although natriuretic peptides function primarily to regulate electrolytes, they are expressed in immune cells and there is increasing evidence of their role in inflammation and immunity^38^.

### Immune cell subtype analysis identifies M1 macrophages as strong correlate of ICB response

To identify immune cell populations associated with ICB response, we estimated the immune cell composition of each tumor using Cibersortx^39^ and compared the abundance of immune cell subsets between ICB responders and non-responders. M1 macrophages, T-follicular helper (Tfh), and activated CD4 memory T-cells had higher abundance in ICB responders as compared to non-responders (**Figure 3A, Supplementary Figure 3**). M1 macrophages are known to be pro-inflammatory and anti-tumoral^40^. A recent study found M1 macrophage infiltration to be predictive to ICB response in urothelial cancer^41^. Our results further demonstrate that M1 macrophage infiltration could be a general feature of ICB response across multiple cancer types. Tfh cells are specialized CD4+ T cells that aid the formation of germinal centers in tertiary lymphoid structures (TLS)^42^, interacting with B-cells to activate antibody responses^43^ and enhancing CD8+ T-cell effector functions^44^. While the presence of Tfh cells has been associated with favorable survival in multiple cancer types^30,43–45^, their putative role in ICB response has not been studied. Our analysis also demonstrated that while resting CD4 memory T-cells were enriched in non-responders, activated CD4 memory T-cells were enriched in ICB responders. This suggests that pre-existing CD4+ T-cell immunity is important for effective anti-tumoral response upon ICB treatment.

**Figure 3.**
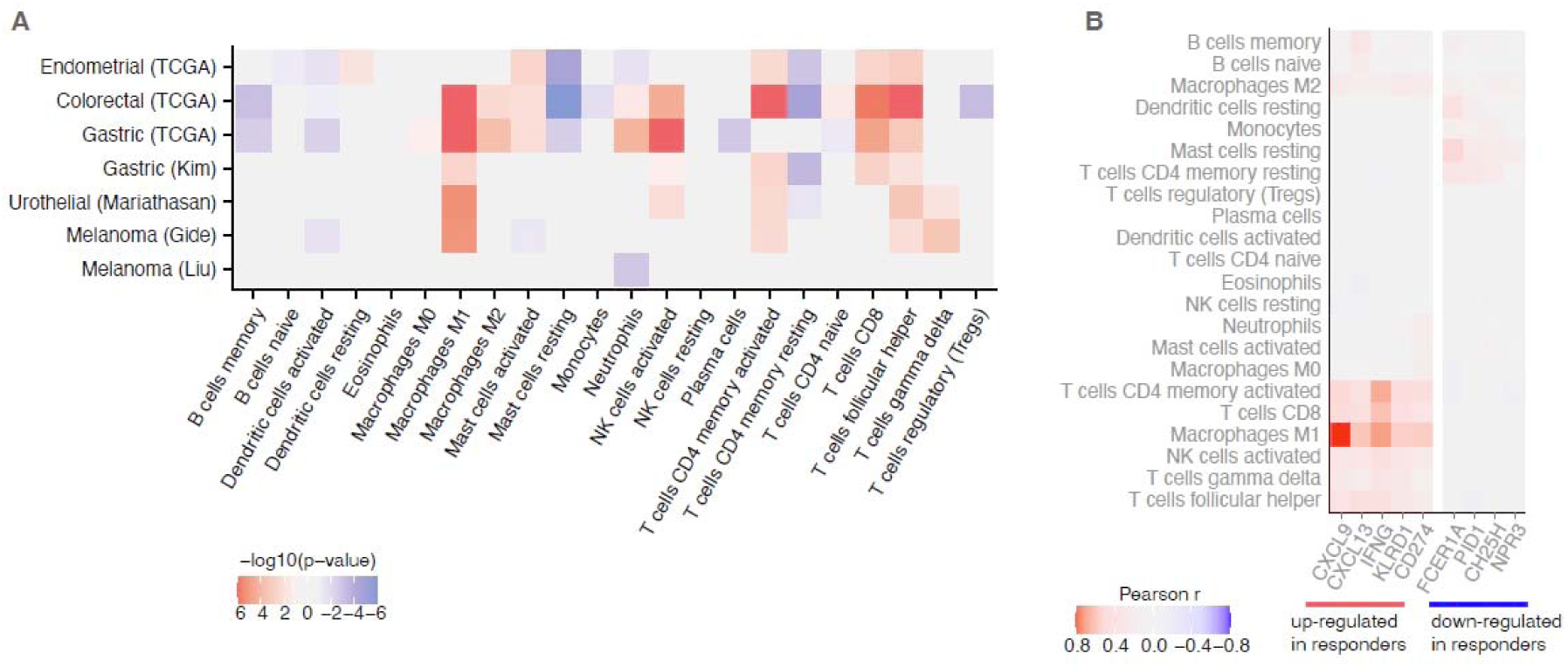
Immune cell type enrichment analysis. Abundance of immune cell types of each tumor is estimated using Cibersortx. **(A)** Heatmap of cell types analyzed. Heatmap is colored by the signed -log10(p-value) of the difference in Cibersortx scores between responders (MSI) and non-responders (MSS). **(B)** Correlations between the abundance of immune cell types with the gene expression of top stromal biomarkers.

Next, we examined associations between the top DEGs and infiltration of specific immune cell subsets. We found that the top gene expression markers of good ICB response were generally associated with abundance of M1 macrophages, CD4+/CD8+ T-lymphocytes, and activated NK cells (**Figure 3B, Supplementary Figure 4**). Using orthogonal expression data of immune cells subtypes from flow cytometry and single-cell RNA-seq experiments, we confirmed that *CXCL9* expression was strongly associated with M1 macrophage infiltration, while *CXCL13* and *IFNG* expression were strongly associated with T-cell infiltration (**Supplementary Figure 5**). Interestingly, the DEGs associated with poor ICB response, including FCER1A, were correlated with resting state of mast cells, CD4+ memory T cells, and dendritic cells, suggesting that these genes are likely markers of a quiescent or immune-suppressive microenvironment (**Figure 3B, Supplementary Figure 4**).

### Cancer intrinsic features of ICB response tend to be tumor type specific

Previous studies have proposed multiple mechanisms by which cancer cells may directly suppress tumor immunity and impede ICB therapy^16,46^. Surprisingly, our meta-analysis across tumor types revealed a paucity of DEGs associated with ICB response in cancer cells. In cancer cells, we identified 9 genes differentially expressed between ICB responders and ICB non-responders across tumor types (**Figure 4A, Supplementary Figure 2B & D**). In addition, we observed that the differential expression signal for these 9 genes was mostly driven by the MSI ICB-responder-enriched cohorts. As compared to stromal DEGs (**Figure 2A**), the signal for the cancer DEGs was less consistent across the distinct ICB cohorts (**Figure 4A**). Therefore, we hypothesize that the cancer DEGs identified by the meta-analysis are likely related to biology of MSI tumors, rather than general mechanisms of ICB response. We also confirmed that the paucity of cancer cell DEGs, as compared to stroma cell DEGs, persisted regardless of the significance cutoff in our meta-analysis (**Figure 4B**), and we were not able to identify functional enrichment or relationships among cancer DEGs.

**Figure 4.**
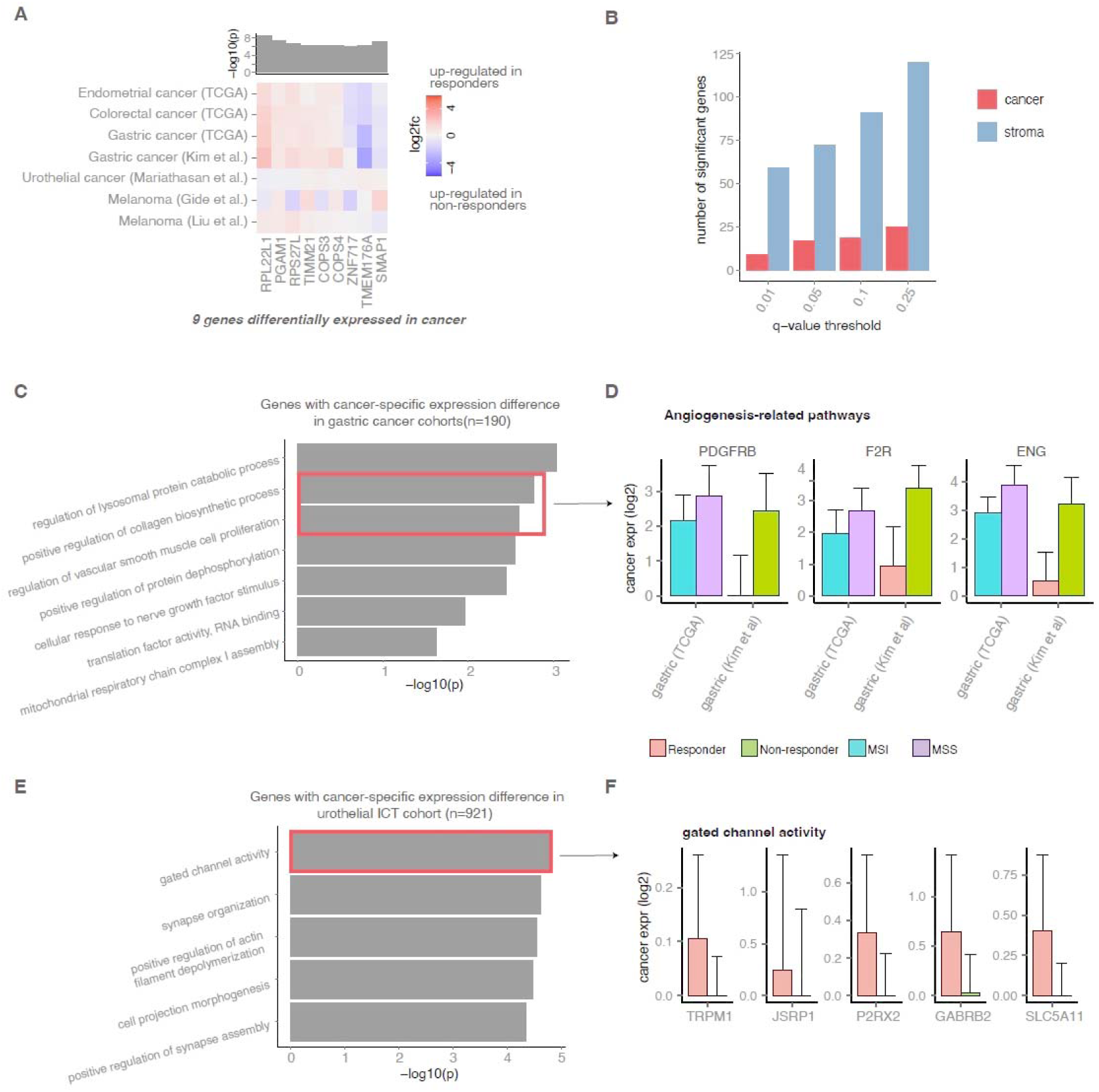
Differentially expressed genes between responders and non-responders in the cancer compartment. **(A)** Heatmap of genes consistently differentially expressed across discovery cohorts in the cancer cells. Heatmaps are colored by the log2 fold change of gene expression. *P*-values of individual cohorts are combined using the Fisher’s method, and the - log10(meta p-value) is presented in the bar plot. Genes with q-value<0.01 and *p*-value<0.1 at least 1 ICB cohort are shown. **(B)** Bar plot shows the number of significant genes found in the stroma and cancer compartment at different q-value cutoffs. Pathway analysis of genes with cancer-specific expression differences in **(C)** gastric cancer and **(E)** urothelial cancer. Bar plots of deconvoluted cancer expression of differentially expressed genes in the top enriched pathway in **(C)** gastric cancer and **(E)** urothelial cancer. Error bars represent estimated standard error.

Next, we conducted the same DEG analysis within individual tumor types to explore the potential for tissue-dependent processes involved in ICB response. For melanoma, we only identified 22 cancer DEGs shared between the 2 melanoma cohorts, and these genes were not enriched in specific pathways. In gastric cancer, we identified 190 cancer ICB response DEGs across the gastric ICB cohort and TCGA cohort. Interestingly, genes associated with poor ICB response were enriched for angiogenesis-related functions (**Figure 4C**), with pro-angiogenic genes ENG, F2R and PDGFRB overexpressed in ICB non-responders (MSI/EBV) compared to responders (MSS) (**Figure 4D**). These data are consistent with the understanding that angiogenesis can be associated with immune suppression^47^, suggesting that a pro-angiogenic environment may impede ICB response in gastric cancer. In urothelial cancer we found that cancer DEGs were enriched for neuron-related functional terms (**Figure 4E**), with neuronal genes overexpressed in ICB responders compared to ICB non-responders (**Figure 4F**). This observation is concordant with a previous study linking the neuronal expression subtype^48^ of urothelial cancer with poor overall survival but high response rate to ICB^49^.

Overall, while these results indicate an overall paucity of tissue-agnostic cancer-cell-intrinsic features of ICB response, they also highlight the potential for tissue and tumor-type specific expression signatures in cancer cells that may predict ICB response.

### Lack of cancer intrinsic immune checkpoint regulation across cancer types

To further investigate if specific ligand-receptor interactions between cancer and immune cells may promote primary ICB resistance, we systematically investigated the expression of 30 known immune checkpoint ligand-receptor pairs in both the cancer and stromal components of the TME (**Figure 5A**). We found stromal overexpression of 13 checkpoint receptors and 2 checkpoint ligands to be associated with ICB response across cancer types. Checkpoint ligands PD-L1 and PD-L2 and their receptor PD-1 were coordinately upregulated in the stroma of ICB responders compared to ICB non-responders. PD-L1 (*CD274*) and PD-1 were significantly upregulated in the stroma of ICB responders in 6 out of 7 cohorts studied (p-value<0.05, permutation test), while PD-L2 was upregulated in 5 out of the 7 cohorts (**Supplementary Figure 6**). Interestingly, no checkpoint ligands were consistently differentially expressed and associated with ICB response in cancer cells across tumor types.

**Figure 5.**
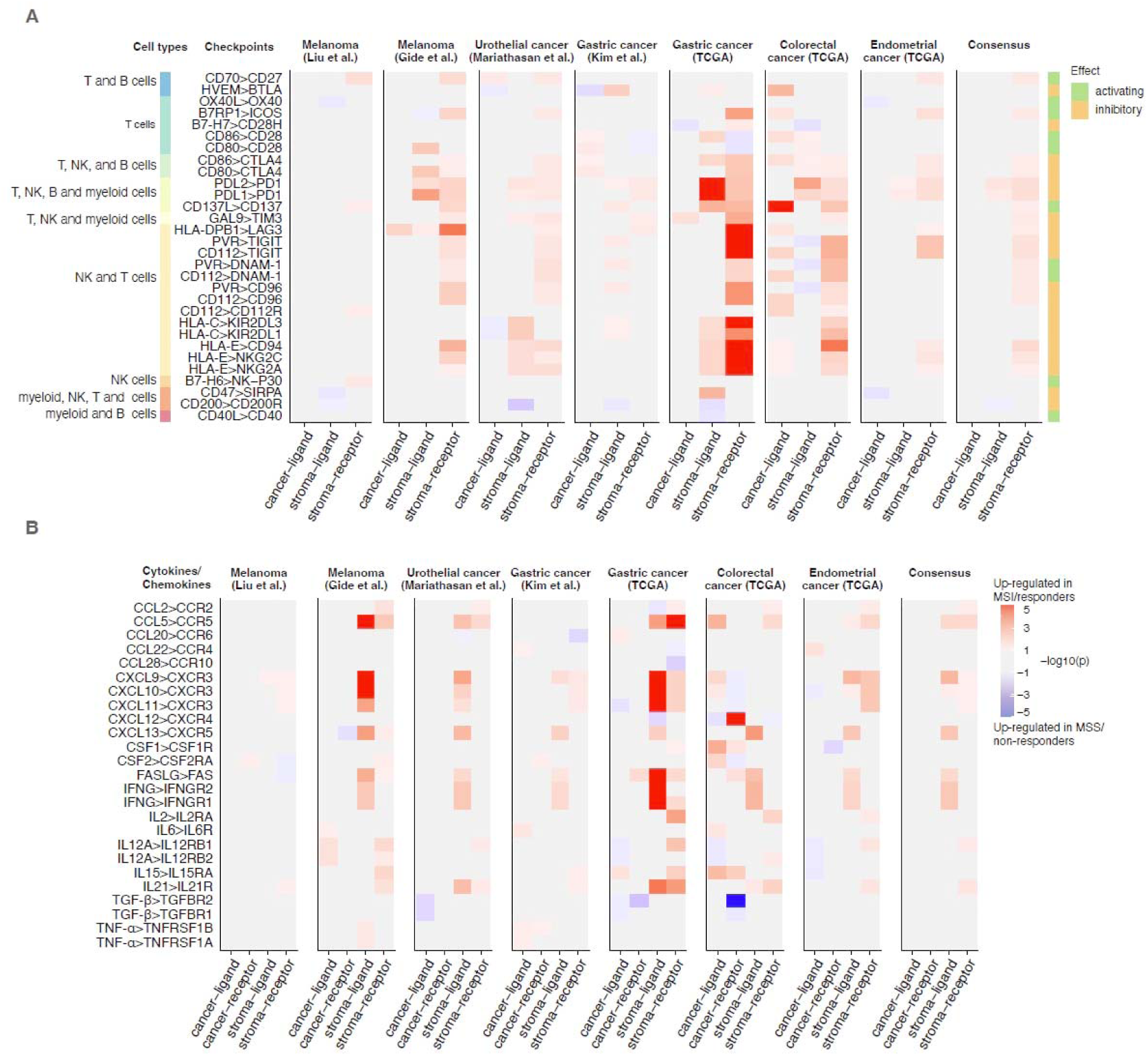
Analysis of ligand and receptor expression of immune modulators. Heatmaps of the inferred ligand and receptor expression in the cancer and stroma compartments for **(A)** immune checkpoint proteins and **(B)** cytokines. Heatmaps are colored by the signed - log10(*p*-value) of differential expression.

In addition to immune checkpoints, cytokines also play important roles in tumor immunity^50–52^. We examined the expression of 13 pairs of cytokines and cytokine receptors known to be involved in tumor immunity^50,52^, aiming to identify cytokine signals associated with ICB response (**Figure 5B**). Similar to our analysis for immune checkpoints, we found that cancer-intrinsic cytokine expression was not associated with ICB response across tumor types. However, six cytokines (*IFNG*, *FASLG*, *CXCL13*, *CCL5*, *CXCL10* and *CXCL9*), and the corresponding receptors for three of the cytokines (*CCR5* for *CCL5*, *CXCR3* for *CXCL9* and *CXCL10*) were significantly upregulated in the stroma of ICB responders, consistent with previous observations^11,25^. Overall, our analysis finds no evidence for a universal mechanism by which cancer cells inhibit immune cells via checkpoints interactions or chemokine/cytokine signaling.

### A 3-feature classifier robustly predicts ICB response across cancer types

Next, we explored if we could build an accurate predictor of ICB response based on the consensus DEGs identified earlier (**Figure 6A**). We focused on the 59 stroma-specific DEGs (**Figure 2A**) as they were more consistent across tumor types, in comparison to cancer DEGs, and therefore more likely to be generalize to unseen cancer types. While a stroma transcriptomic signature would likely reflect the state and favorability of the tumor microenvironment for ICB response, TMB is likely a proxy of the tumor immunogenicity. Since the stroma DEGs and TMB likely represent different facets of tumor immunity, we hypothesized that these could be orthogonal predictors of ICB response and therefore considered both factors for the predictive model. To identify the best predictive features of ICB response, we used LASSO logistic regression with 10-fold cross-validation to select robust features from the set of 59 stromal DEGs and TMB (**Figure 6A**). The final model was built using the top-3 features: TMB, *IFNG* expression, and *FCER1A* expression (**Figure 6B**). The model was trained on 461 samples from the ICB discovery cohorts with both transcriptomic and genomic data. Next, we evaluated the model performance on 2 independent test cohorts. Test cohort 1 consisted of 49 tumors from melanoma patients treated with the PD-L1 inhibitor nivolumab^53^. Test cohort 2 consisted of 87 tumors from 20 solid cancer types treated with a mix of ICB agents^54^. We calculated the area under the receiver operating curve (AUC) of our model, and benchmarked it against established biomarkers of ICB response, namely CXCL9+TMB^14^, T-cell GEP^11^, TIDE^13,55^, TMB alone^2,8^, and PD-L1 expression^3^. Intriguingly, our compact 3-feature predictor outperformed all existing biomarkers. On test cohort 1, the 3-feature model achieved an AUC of 0.66, compared to 0.62 for CXCL9+TMB, 0.64 for T-cell GEP, 0.44 for TIDE, 0.55 for PD-L1 expression, and 0.57 for the TMB-only model. On test cohort 2, the model achieved an AUC of 0.72, compared to 0.67 for CXCL9+TMB, 0.63 for T-cell GEP, 0.32 for TIDE, 0.61 for PD-L1 expression, and 0.55 for the TMB-only model (**Figure 6C**). This demonstrates that *FCER1A,* a novel negative biomarker of ICB response identified from our transcriptome-wide analysis, contributes additional predictive power of ICB response over existing biomarkers. Furthermore, in cohorts that included survival data, model predictions were significantly associated with progression free survival (**Figure 6D**) and, to a lesser extent, overall survival (**Supplementary Figure 7**).

**Figure 6.**
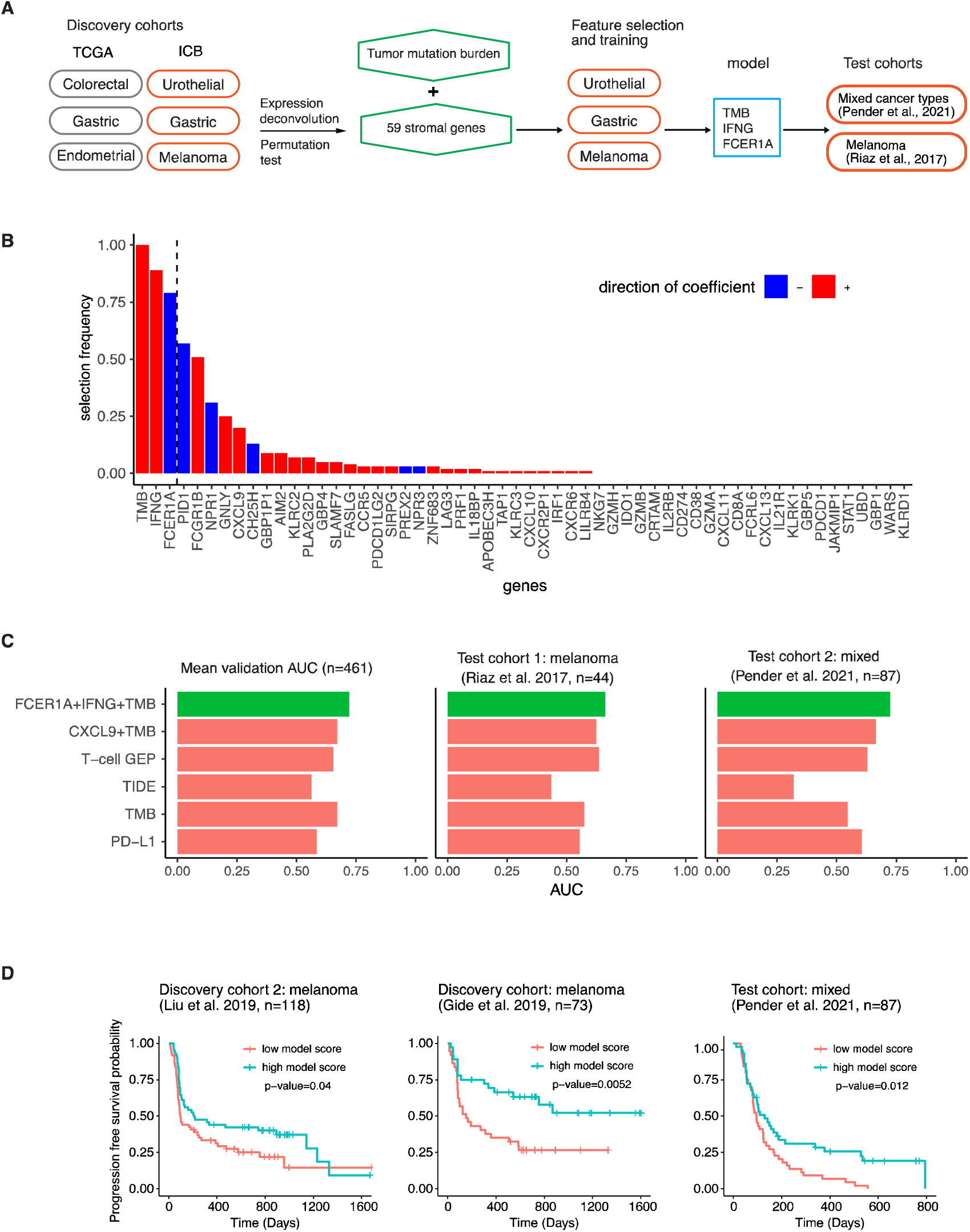
Predictive modeling of response to immune checkpoint inhibition. **(A)** Schematic of data and methods used in model training and testing. **(B)** Top ranked features of ICB response selected by LASSO logistic regression. **(C)** Comparisons of the performance of the TMB+*IFNG*+*FCER1A* model and existing biomarkers in 2 independent test cohorts. **(D)** Kaplan-Meier curves of progression free survival for patients with high vs low predicted scores (greater than or less than median); p-values from log-rank test shown.

## Discussion

In this study, we performed a transcriptome-wide meta-analysis of ICB response in 1486 tumors. These tumor samples included 534 tumors with annotated clinical response to ICB treatment, and 952 tumors from cancer types enriched for MSI and ICB-responding tumors. Using *in silico* tumor transcriptome deconvolution, we analyzed gene expression signatures of cancer and stromal cells and identified differentially expressed genes associated with ICB response across tumor types. Strikingly, this meta-analysis revealed a paucity of ICB-response DEGs in cancer cells as compared to stromal cells. The few cancer DEGs that we identified by the meta-analysis were more likely related to the biology of MSI tumors and their DNA mismatch repair deficiency rather than being general determinants of ICB response. In contrast, our analysis revealed 59 stroma-specific ICB-response DEGs, which were strongly enriched in immune-related pathways and processes (**Figure 7**).

**Figure 7.**
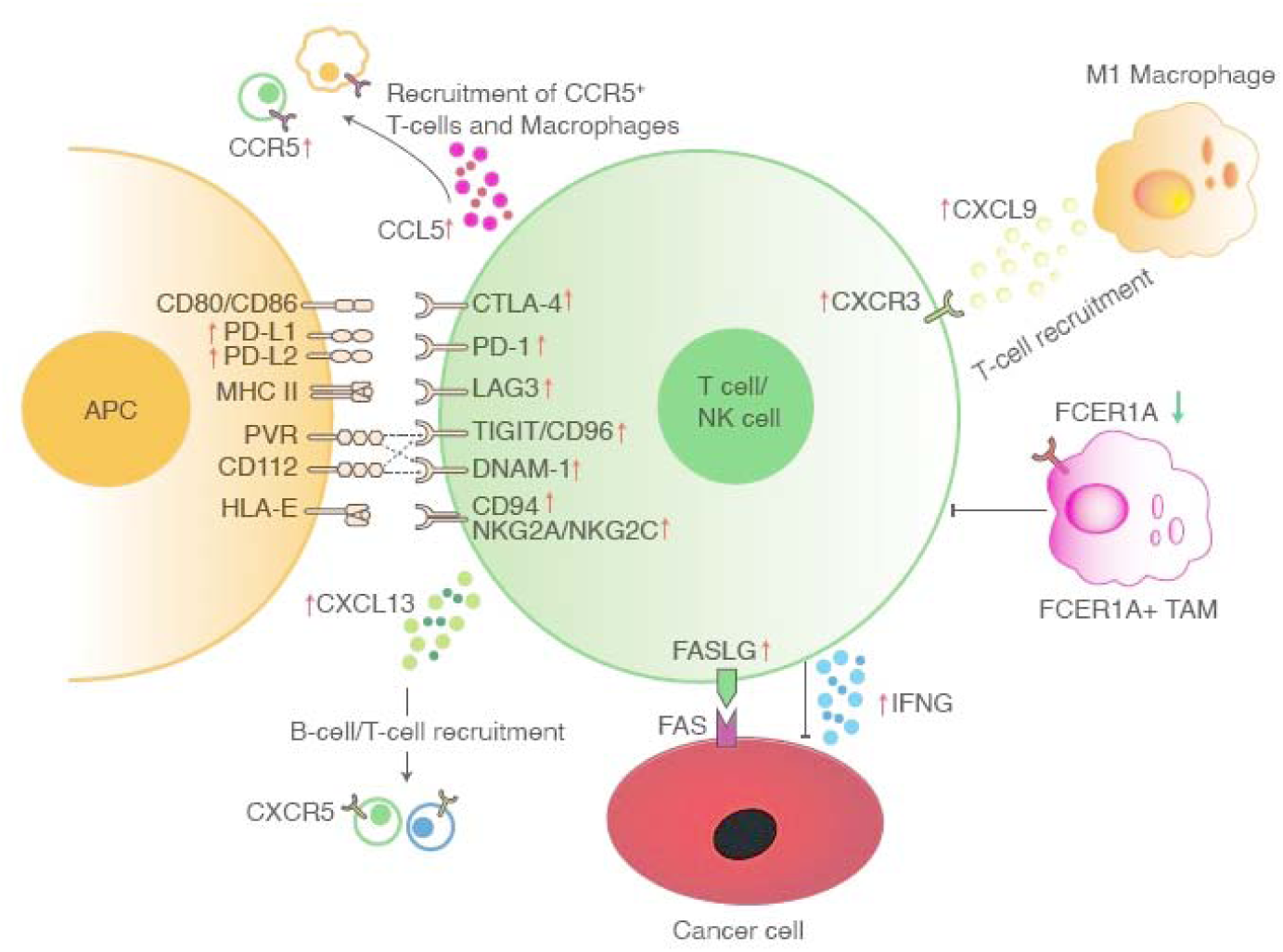
Schematic summarizing the key components associated with ICB response. Red arrows indicate gene positively correlated with ICB response. Green arrow indicates gene negatively correlated with ICB response.

The two chemokine ligands, *CXCL9* and *CXCL13*, constituted the top-2 stromal genes overexpressed in ICB responders. Underlining the validity of our approach, these two genes have been highlighted for their roles in anti-tumor immunity and association with favorable ICB response in multiple recent studies^14,31,32^. Previous studies suggest CXCL13 may play dual roles in ICB response. First, CXCL13 is produced by Tfh cells and involved in B-cell organization in TLS ^29^, and the presence of TLS can be a positive predictor of ICB response^30,44^. Second, *CXCL13* has been reported to be highly expressed by neo-antigen reactive CD8+ T-cells directly involved in cancer cell killing ^56,57^. In line with these findings, we found enrichment of both Tfh and CD8+ T-cells in tumors of ICB responders compared to non-responders across cancer types (**Figure 3,7**). Additionally, the abundance of these two cell types correlates with *CXCL13* expression. In agreement with recent work, we found that *CXCL9* expression is strongly correlated with M1 macrophage infiltration^31,58^, and M1 macrophages were one of the most differentially enriched immune cell types in ICB responders (**Figure 3a**). Although myeloid infiltration has traditionally been associated with immune suppression^40^, our data is consistent with the hypothesis that M1 macrophage infiltration, and macrophage polarization, could be an important determinant of tumor sensitivity to ICB.

Intriguingly, we also identified a set of immune-related genes that were negatively associated with ICB response. Among these genes, *FCER1A* and *CH25H* have previously been reported to be over-expressed in anti-inflammatory M2 macrophages^35,59^, and FCER1A+ TAMs have been reported to promote tumor progression by engaging in a positive feedback loop with tumor initiating cells^36^.

Interestingly, we observed an enrichment of the resting states of CD4 memory T-cells, dendritic cells and mast cells in ICB non-responders. In contrast, the activated states of these cell types are enriched in the tumors of ICB responders (**Figure 3B**). Furthermore, while *FCER1A* is expressed in antigen presenting cells like dendritic cells and macrophages, we noted that its expression is 25 times higher in resting dendritic cells compared to activated dendritic cells, and 14 times higher in M2 macrophages compared to M1 macrophages^39^. These results suggest that a pre-existing immunosuppressive microenvironment, associated with resting immune cells and high *FCER1A* expression, may prevent effective anti-tumor immune responses in ICB non-responders.

Analyses of immune checkpoint and cytokine ligand-receptor interactions revealed coordinated upregulation of several ligand-receptor pairs in the stroma of tumors from ICB responders, including genes encoding PD-L1/PD-L2 and their receptor PD-1, CXCL9/CXCL10 and their receptor CXCR3, and CCL5 and its receptor CCR5 (**Figure 5**, **Figure 7**). In contrast, we did not find consistent expression changes of checkpoint proteins and cytokines in cancer cells of ICB responders. Among chemokines upregulated in ICB responders, CXCL9, CXCL10, and CXCL13 promote lymphocyte recruitment, while CCL5 is a chemoattractant of myeloid cells. CCL5 recruits CCR5+ blood monocytes and neutrophils that can differentiate into immunosuppressive tumor associated macrophages and neutrophils^50,51^. In addition, CCL5 also has direct pro-tumoral effects on cancer cells, promoting cancer proliferation and metastasis^50,51^. Preclinical and clinical studies suggested that targeting the CCL5-CCR5 axis could enhance anti-tumoral response^60,61^. As *CCL5* and *CCR5* are both overexpressed in ICB responders, our result suggests that targeting CCL5-CCR5 signaling in conjunction with ICB might further potentiate anti-tumoral immune response. Currently, the majority of companion diagnostics for PD-1/PD-L1 inhibitors measure PD-L1 expression level on either cancer cells alone, or on both cancer cells and infiltrating immune cells, but seldomly on immune cells only^16,62^. Our study strongly suggests that PD-L1 expression on immune cells is a more consistent determinant of ICB response across tumor types.

Using the stromal consensus DEGs identified from the meta-analysis, we developed a model to predict ICB response from gene expression data and TMB. This 3-feature model comprised of a single positive marker of immune infiltration (*IFNG*), a single negative marker of immune suppression (*FCER1A*), in combination with TMB. The model robustly predicted ICB response across both the discovery cohort and 2 independent test cohorts, outperforming existing gene expression signatures for ICB response. While ideal pan-cancer biomarkers of ICB response should be robust across patient cohorts, we noticed that the Liu et al. melanoma cohort did not share many DEGs with the other cohorts, including other melanoma cohorts. This result highlights significant heterogeneity between ICB cohorts, even among cohorts of the same tumor type. It is unclear to what extent this heterogeneity arise from patient selection, differences in ICB regime, differences in prior treatments, different sample collection protocols, or differences in gene expression assays. This highlights the need for large, consistently selected patient cohorts, with standardized gene expression profiling to further improve biomarker discovery.

As cancer cells are under negative selective pressure from the immune system, several mechanisms of immune escape by cancer cells have been proposed: (i) cancer cells down-regulate antigen presentation by deleting HLA alleles^46^, (ii) cancer cells deplete neoantigens to avoid detection^63^, (iii) oncogenic signaling may up-regulate PD-L1 expression and suppress T-cell recruitment^64^. However, the prevalence of these immune evasion mechanisms are still unclear. A recent meta-analysis found no association between loss of heterozygosity of HLA alleles and ICB response across ICB cohorts^14^. Furthermore, a pan-cancer analysis failed to detect neo-antigen depletion signals in most tumor types^65^. Similarly, we found a lack of conserved cancer-intrinsic mechanisms of immune evasion on the transcriptomic level, which challenge the prevailing dogma that cancer cell-intrinsic expression of checkpoint ligands can modulate immune activity and predict ICB response across cancer types. These insights have profound implications on biomarker assays and *in vitro* model systems for development of improved ICB drugs. These results suggest that cancer cells adopt diverse mechanisms of immune evasion, with no universal strategy adopted by cancer cells to avoid immune-mediated killing. The growing number of molecularly characterized ICB cohorts will enable further exploration of the importance and prevalence of the different cancer-intrinsic immune evasion strategies in determining ICB response.

## Methods

### Clinical cohorts treated with ICB

Data from the following studies were used for discovery:

1. Kim et al.^22^ cohort comprises of 45 metastatic gastric cancer patients treated with PD-1 inhibitor pembrolizumab
2. Mariathasan et al.^23^ cohort comprises of 298 metastatic urothelial cancer patients treated with PD-L1 inhibitor atezolizumab
3. Liu et al. ^25^ comprises of 118 metastatic melanoma patients treated with PD-1 inhibitors nivolumab or pembrolizumab
4. Gide et al. ^24^ comprises of 73 metastatic melanoma patients treated with anti-PD-1 alone (nivolumab or pembrolizumab, or with anti-CTLA-4 (nivolumab or pembrolizumab with ipilimumab)

Data from the following studies were used for validation:

1. Riaz et al. ^53^ comprises 49 advanced melanoma patients treated with PD-L1 inhibitor nivolumab.
2. Pender et al.^54^ comprises of 87 metastatic tumors from 20 solid cancer types treated with a mix of ICB agents.

### Gene expression data collection and normalization

Expression data of TCGA colorectal (COAD, READ), gastric (STAD) and endometrial (UCEC) tumors were downloaded from the UCSC Xena repository. Raw genomic and transcriptomic data of Kim et al.^22^ gastric ICB cohort were downloaded from the European Nucleotide Archive, and processed using the bcbio nextgen pipeline. Briefly, RNAseq reads were aligned to the human transcriptome using STAR^66^, and TPM counts were quantified using SALMON^67^.

Processed transcriptomic data of Mariathasan et al. (urothelial)^23^, Liu et al. (melanoma)^25^, Gide et al. (melanoma)^24^, Riaz et al. (melanoma)^53^, and Pender et al. (mixed)^54^ ICB cohorts were obtained from the original publications. Genes with 0 expression in more than 50% of the samples were removed. All gene expression data were log2 transformed and upper quantile normalized across all cohorts.

### Tumor purity estimation

Purity estimates of TCGA tumors were obtained from a previous publication^20^, where consensus purity was estimated from mutation allele frequencies, copy number variations and RNA profiles of each tumor. Tumor purity of samples from ICB cohorts were estimated using PUREE^68^, a machine learning method for tumor purity estimation from bulk RNA transcriptomics. For the Kim et al. gastric ICB cohort^49^ where genomic data is also available, we calculated consensus purity estimates based on both genomic and transcriptomic data. We first computed purity estimates using 2 RNA-based methods (PUREE^68^ and Estimate^69^), and 2 DNA-based methods (Sequenza^70^, PurBayes^71^). Then, we performed quantile-quantile normalization on the purity estimates of individual methods, and took the mean normalized purity estimate as the consensus purity.

### MSI status and ICB response classification

MSI status of TCGA tumors were obtained from two previous pan-cancer studies of microsatellite instability^9,10^. EBV status of the TCGA STAD tumors was obtained from a previous TCGA study on gastrointestinal cancers^72^.Clinical annotations of ICB response based on the RECIST criteria were obtained from the original studies. Consistent with recent literature^14,23^, we classified “CR/PR” as responders and “SD/PD” as non-responders.

### Expression deconvolution

We modelled bulk tumor expression as the sum of cancer and stroma expression, weighted by cancer purity^20^. This can be written as,

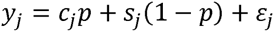

where *y_j_* is a *n* × 1 vector of bulk tumor expression for gene *j* in *n* samples, *c_j_* and *s_j_* are the average cancer- and stroma-specific expression for a gene *j*, *p* is *n* × 1 vector of cancer purity, and *ε_j_* is the residual. Given bulk expression (*y_j_*), and tumor purity (*p*) non-negative least squares (NNLS) regression can be used to estimate cancer- (*c_j_*) and stroma-specific (*s_j_*) expression for each gene *c_j_* and *s_j_* represent estimates of mean cancer- and stroma-specific expression for a gene *j* for *p_c_* = 1 and *p_s_* = 0, respectively. The standard error of these predicted values for gene *j* is calculated as,

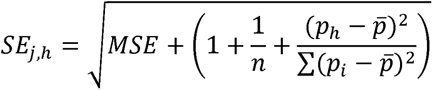

where *p_h_* = *p_c_* = 1 for cancer-specific expression and *p_h_* = *p_s_* = 1 for stroma-specific expression; *MSE* is the mean square error.

### Permutation statistic to identify differentially expressed genes

To test differences in cancer (or stroma) expression between two response groups (e.g. *C_Rj_* - *C_NRj_* for gene *j*), the above approach is applied to each group separately, and a test statistic is calculated as,

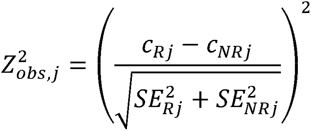

To assess significance under the null hypothesis of no difference in cancer (stroma) expression between the 2 response groups, a null distribution is generated by permutation, where group labels are first randomly shuffled 10,000 times and the above procedure is applied to each permuted dataset, generating a null distribution of 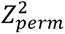 statistics. A p-value is calculated as,

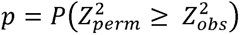

That is, the probability of observing 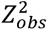 or larger, given the null hypothesis is true.

### Identification of consensus DEGs across cancer types

For each gene and each tumor compartment (caner or stroma), a meta p-value is computed by combining the permutation p-values of all cancer cohorts using Fisher’s method. We used the Bonferroni correction to adjust for multiple testing to maintain the overall α at 0.01. Genes that are differentially expressed in different directions in at least 2 cohorts were removed. Finally, to eliminate DEGs that are involved in MSI biology but are not directly associated with ICB response, we require the final list of consensus DEGs to be differentially expressed in at least 1 ICB cohort (nominal p-value<0.1).

### Pathway enrichment analysis

To find out if the DEGs identified are enriched in specific biological processes, pathway enrichment analysis was performed using ClueGo^73^ with default parameters. Briefly, two-sided-hypergeometric statistic was used to compute the enrichment of a given list of DEGs in GO biological processes terms, and the resulting p-values were corrected for multiple testing using the Bonferroni step down approach. To aid interpretation, significant biological terms with high overlap as measured by the kappa coefficient were merged into functional groups. To calculate the enrichment of stroma DEGs in immune-related functions, we defined immune-related genes as genes in the “immune system process” term and its child terms in the GO biological processes dataset. Additionally, we manually curated stroma DEGs not labeled as immune genes by GO, and further labeled *FCER1A*, *GZMH*, *FCRL6*, *SIRPG*, *JAKMIP1*, and *WARS* as immune-related genes based on literature search.

### Building a predictive model of ICB response

We used 461 patients from 3 discover cohorts (Mariathasan et al^23^, Liu et al^25^, Kim et al.^22^) with both transcriptomic data and TMB as training data. TMB was obtained from the original publications and standardized into Z-scores (mean=0, standard deviation=1). To identify the most predictive features of ICB response, we used the least absolute shrinkage and selection operator (LASSO) to select features from the 59 stroma DEGs identified earlier and TMB. LASSO penalizes the sum of the absolute values of the regression coefficients, shrinking some coefficients to zero, and thereby identifies a simpler and more interpretable model. The regularization parameter λ was chosen by 10-fold cross-validation such that the error of the selected model was within 1 standard deviation from the minimum error. LASSO regression and cross validation were performed using the ‘glmnet’ package in R. To select for the most robust features, we bootstrapped 100 samples with 90% of the data in each bootstrap, and selected features that are found by at least 75% of the bootstraps for the final model. The final logistic regression model is built on all 461 training samples using the features selected by LASSO (TMB+IFNG+FCER1A).

### Benchmarking ICB prediction model against existing biomarkers

T-cell GEP scores were calculated from normalized RNAseq data using the weights provided by Ayers et al. 2018^11^. TIDE scores were calculated from normalized RNAseq data using the TIDE web platform^55^. The CXCL-9+TMB model was trained on our training cohort using logistic regression.

### Immune cell enrichment analysis

Immune deconvolution was performed using CibersortX (with the LM22 signature matrix)^39^ to estimate the absolute abundance of immune cell subsets for each tumor. Immune cell subsets that are differentially infiltrated between responders and non-responders were identified using the Wilcoxon rank-sum test.

## Data and code availability

The code used to generate figures in the manuscript is provided in **Supplementary Data 1**.

## Acknowledgements

This study was supported by the Singapore Ministry of Health’s National Medical Research Council under its OF-IRG program (OFIRG18may-0075) and Singapore Agency for Science, Technology and Research (A*STAR) under its Biomedical Engineering Programme (C221118001 and C221318004)

