## Supplementary Figures for "A transcriptome-wide meta-analysis reveals lack of cancer-cell intrinsic determinants of response to immune checkpoint blockade"

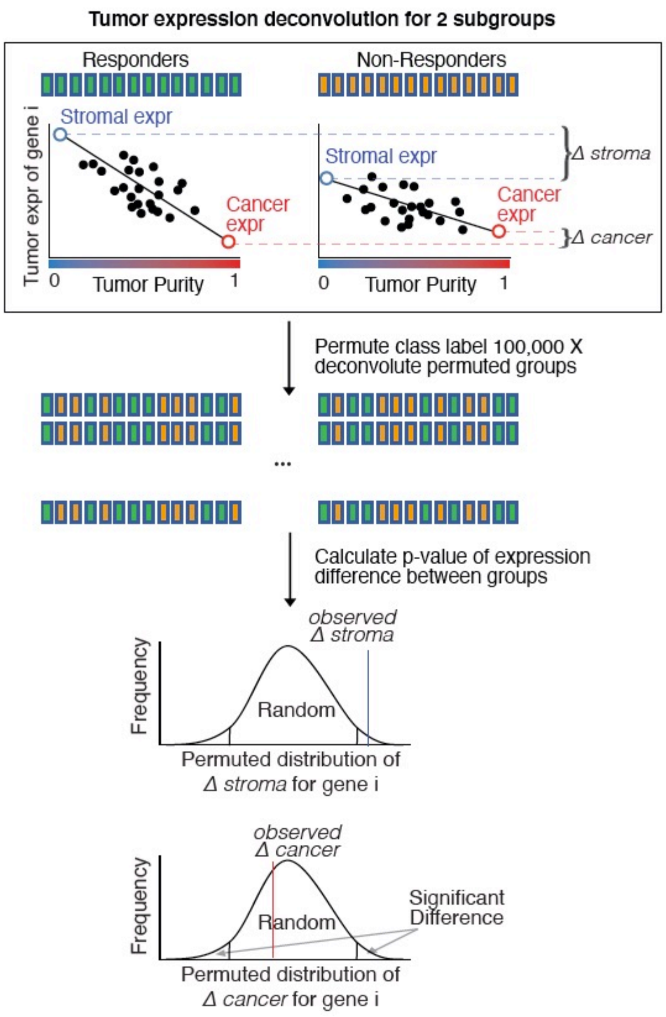


**Supplementary Figure 1**. **Schematic of bulk tumor expression deconvolution and permutation test to identify differentially expressed genes between responders and non-responders in both cancer and stroma compartments.**

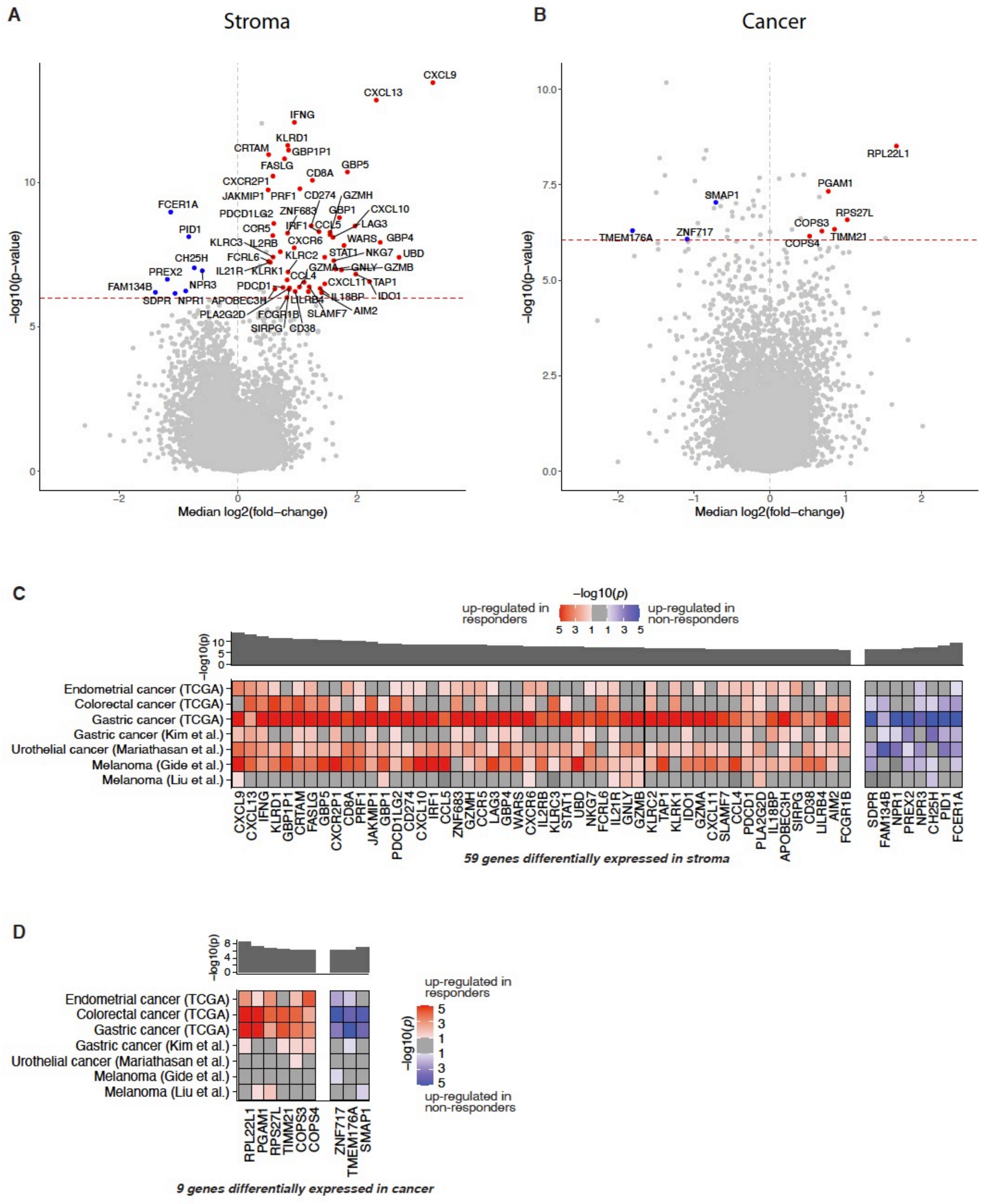


**Supplementary Figure 2**. **Identification of genes that are differentially expressed between responders and non-responders consistently across discovery cohorts.** Volcano plots of differentially expressed genes in the **(A)** stromal and **(B)** cancer compartments. The y-axis is the -log10(meta p-value) of differential expression. The meta p-value is calculated using the Fisher's method, combining p-values all cohorts. The x-axis is the mean log2(fold-change) of gene expression of responders (MSI/EBV) over non-responders (MSS). The red line represents the q-value<0.01 cutoff. Genes with median absolute log2(fold-change)>0.5, q-value <0.01, and p-value<0.1 in at least one ICB cohort are colored and labelled. Heatmaps of differentially expressed genes in the **(C)** stroma and **(D)** cancer compartments. Heatmaps are colored by the signed -log10(*p*-value) of differentially expression. Genes with median absolute log2(fold-change)>0.5, q-value <0.01, and p-value<0.1 in at least one ICB cohort are shown.

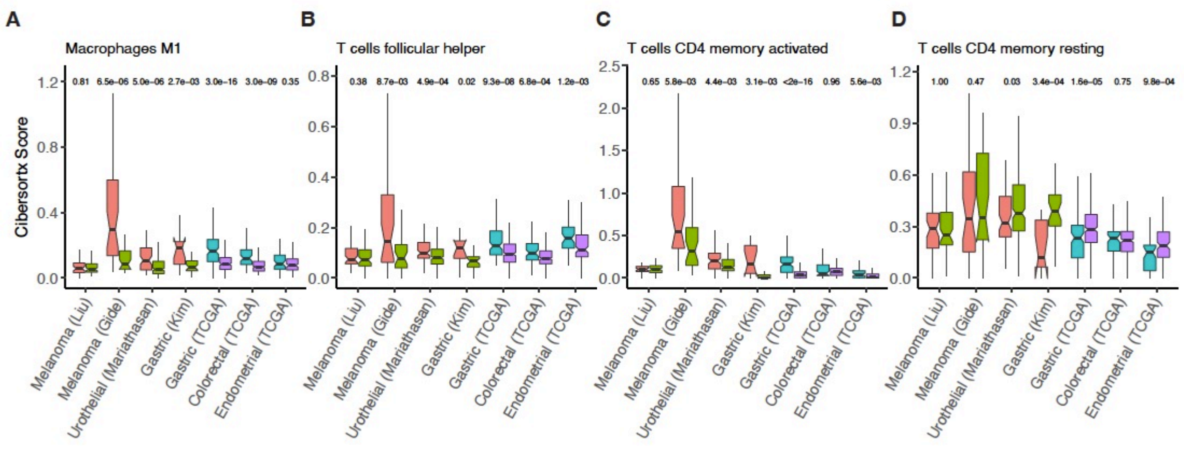


**Supplementary Figure 3**. **Differentially enriched immune cell types between responders and non-responders.** Boxplots showing the Cibersortx scores of 4 cell types that are significantly different between responders (MSI) and non-responders (MSS).

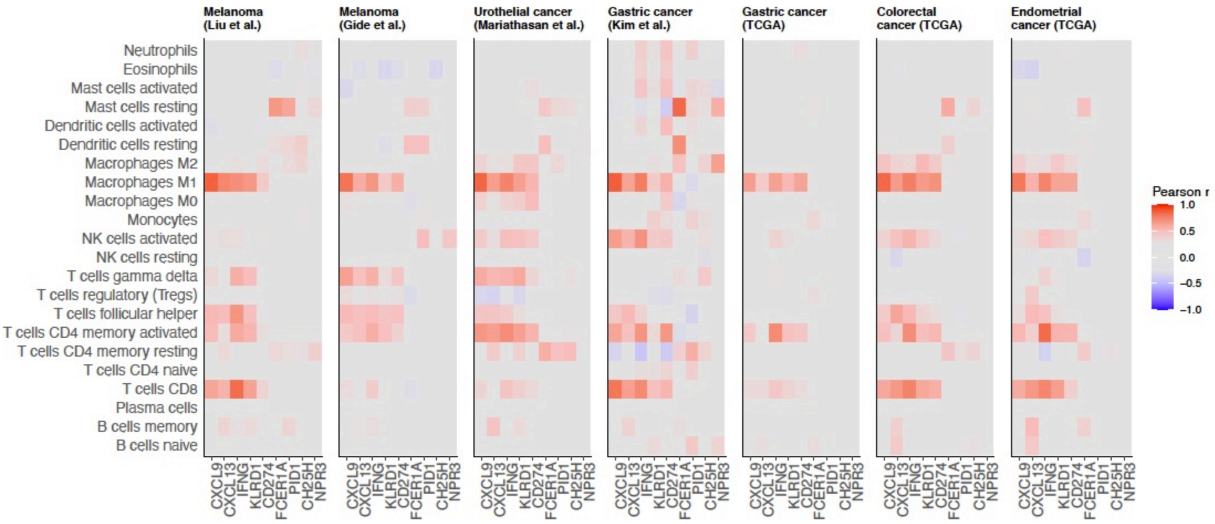


**Supplementary Figure 4**. **Correlations between the abundance of immune cell subtypes with the gene expression of top stromal biomarkers in individual cohorts.**

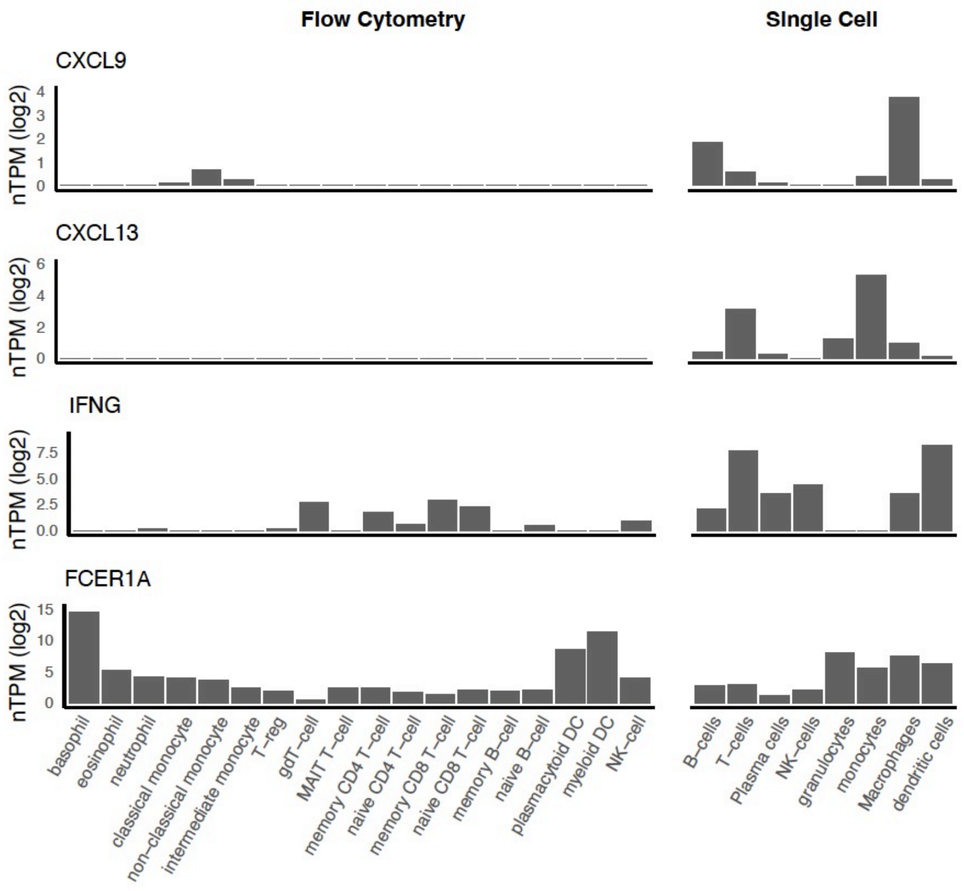


**Supplementary Figure 5**. **Expression of top differentially expressed genes in immune cell subtypes.** Normalized expression of 4 representative stromal biomarkers in different immune cell types. Immune cell expression data is derived from single-cell and flow cytometry sorted RNA-seq experiments from the Human Protein Atlas.

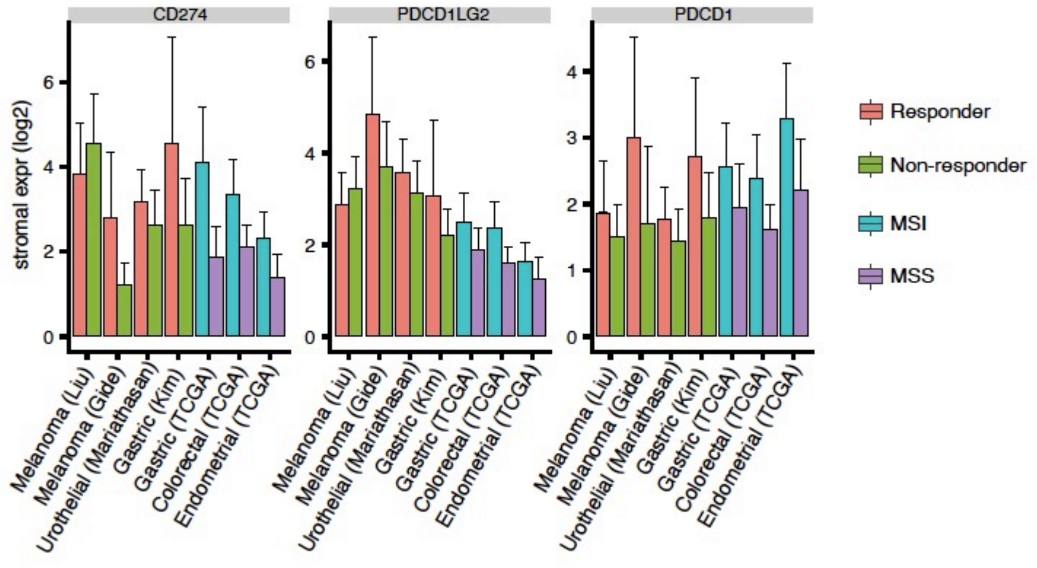


**Supplementary Figure 6**. **Differential expression of the PD1 receptor and its ligands PD-L1 and PD-L2 in the stroma.** Bar plots of deconvoluted stromal expression of the PD1 (*PDCD1*) receptor and its ligands PD-L1 (*CD274*) and PD-L2 (*PDCD1LG2*). Error bars represent estimated standard error.

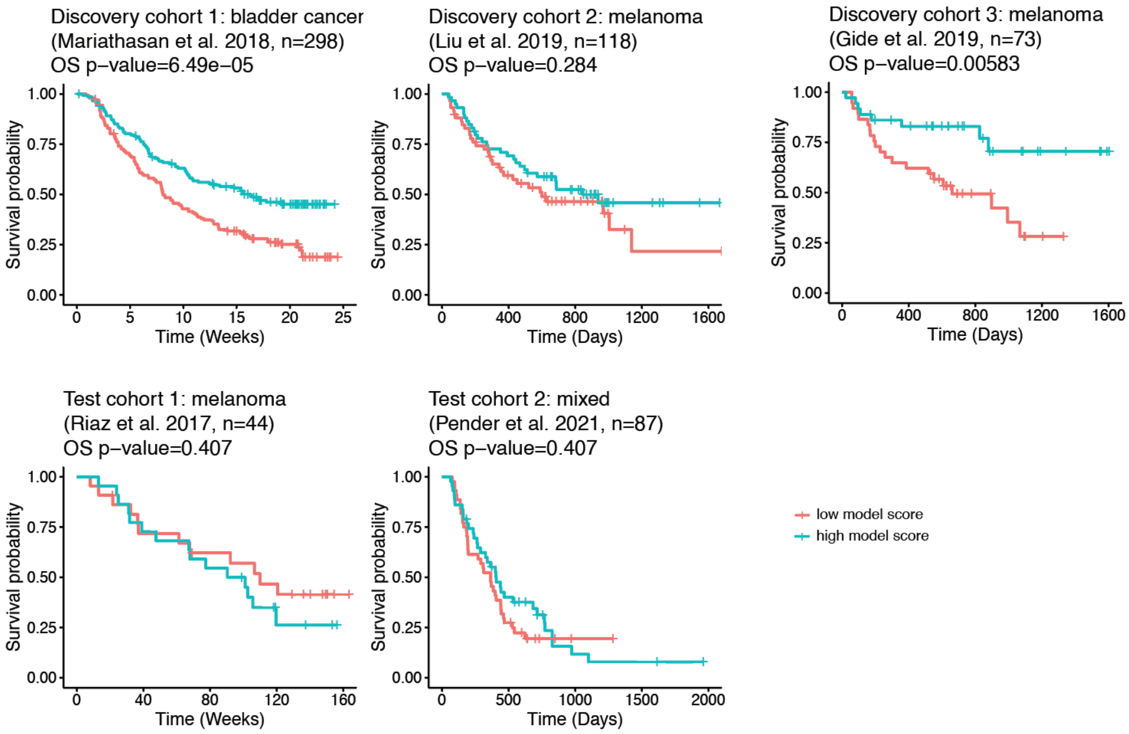


**Supplementary Figure 7**. **Association between model prediction and overall survival.** Kaplan-Meier curves of overall free survival for patients with high vs low predicted scores (greater than or less than median); p-values from log-rank test shown.

| | | | | | | Response | | |
|---|---|---|---|---|---|---|---|---|
| **cohort** | **type** | **PMID** | **N** | **cancer type** | **ICB agent** | **response** | **non-response** | **total** |
| Mariathasan | discovery | 29443960 | 298 | Bladder | atezolizumab | 68 | 230 | 298 |
| Kim | discovery | 30013197 | 45 | Gastric | pembrolizumab | 12 | 33 | 45 |
| Liu | discovery | 31792460 | 118 | Melanoma | nivolumab or pembrolizumab | 47 | 71 | 118 |
| Gide | discovery | 30753825 | 73 | Melanoma | pembrolizumab or nivolumab alone, or with ipilimumab | 40 | 33 | 73 |
| Riaz | validation | 29033130 | 49 | Melanoma | nivolumab | 10 | 39 | 49 |
| Pender | validation | 33020056 | 87 | Lung (20), Melanoma (18), Breast (13), Others (36) | PD-1/PD-L1 (57), CTLA4 (3), combo (26), others (1) | 15 | 72 | 87 |

**Supplementary Table 1. Clinical characteristics of the study cohorts**
